# SLICK: A Sandwich-LIke Culturing Kit for *in situ* Cryo-ET Sample Preparation

**DOI:** 10.1101/2025.02.14.638381

**Authors:** Qiuye Li, Li Zhang, Qin Xu, Peng Zhang, Shiwei Zhu

## Abstract

*In situ* cryo-electron tomography (cryo-ET) has recently been widely used in observing subcellular structures and macromolecules in their native states at high resolution. One of the reasons that it has not been more widely adopted by cell biologists and structural biologists is the difficulties in sample preparation. Here we present the Sandwich-LIke Culturing Kit (SLICK), simplifying the procedure and increasing the throughput for sample preparation for *in situ* cryo-ET (69 words).

## Main

*In situ* cryogenic-electron tomography (*in situ* cryo-ET), combines the advantages of both high-resolution structural biology and *in situ* cell biology^1^, provides a high-resolution view of intracellular space^2^ and can be used to determine the near-atomic resolution structure of abundant macromolecules within the cell^3^. However, electron microscopy (EM) imaging has not been as widely used as light microscopy (LM) in cell biology studies, even with the recent rapid advancement of data collection^4^ and data processing methods^5^. For established users, *in situ* cryo-ET studies still suffers from low throughput. One of the main reasons behind these limitations is the difficulties in sample preparation. To visualize commonly used adhesive mammalian cells by EM, cells need to be cultured on EM grids. The current practice to do so is to place EM grids directly within Petri dishes during culturing^6,7^, which often requires a long training period and still yielded unsatisfied results. Firstly, loosely placed grids tend to move or float during common cell experiment procedures like coating, washing, mixing, or even when moving the Petri dish, potentially damaging grids or cells. Moreover, rearranging misplaced EM grids using tweezers will also cause similar damage. Secondly, it is difficult to pick up a grid from the Petri dish for vitrification, since it is theoretically not possible to grasp the edge of a flat grid from a flat surface. To do so, one usually has to take advantage of small imperfections of grids and dishes, use ultra fine tipped tweezers, and practice for long periods of times. Even so, the EM grids are often bent or scratched in this process, ending up with suboptimal specimens for subsequent EM imaging and wasting the precious samples. Moreover, this step can take the users unpredictably long time, leaving the cells in a unpreferable environment and preventing more time sensitive experiment. With these two difficulties and a lack of dedicated culturing devices, *in situ* cryo-ET studies of adhesive cells are often considered to have a higher technical barrier than LM studies or cryo-ET studies of suspended molecules or cells.

To tackle such problems, we designed a culturing system for *in situ* cryo-ET sample preparation that has three features: 1) EM grids are limited within a designated space with minimal movement; 2) a special bottom surface that allows easy pickup of grids using tweezers; 3) compatible with present cell culturing practice and does not introduce further complications. Furthermore, all the components of this culturing kit are easily accessible and affordable in typical cell biology labs, significantly lowering the technical barrier of *in situ* cryo-ET sample preparation.

This culturing kit, denoted the Sandwich-LIke Culturing Kit (SLICK), contains a 3D-printed plastic disk as the base and a glass coverslip as the lid (**Fig. 1 a**). When culturing, EM grids will be sandwiched between the base and the lid right after glow discharge, and will only be taken out right before vitrification (**Extended Fig. 1**). Therefore, each grid will only be directly handled twice, minimizing the chance of damage (**Extended Fig. 2**). The fully assembled kits with EM grids in them will be handled as a whole. For example, the basic type of SLICK can house 4 EM grids and can be placed in a 35 mm diameter Petri dish (**Fig. 1 b** and **c**). Within the kit, the EM grids are secured in all three dimensions:1) the base has multiple grid slots that are surrounded by three separate pillars, limiting the horizontal movement of the grids (**Fig. 1 d**); 2) the lid is placed 1.5 mm above the grids and semi-locked by a set of lid locks on the base, limiting the floating and flipping of the grids (**Fig1. F** and **Supplementary Video 1**). When fully assembled, the grid will stay in place during normal cell culture operations that involve agitating and mixing (**Supplementary Video 1**). On the SLICK base, an observation hole is placed at the center the grid slots so the cells on grids can be monitored real-time within SLICK (**Fig. 1 e**). We also designed a trench that points at the center of the grid but extends only to the edge of the grids, allowing convenient pick up of the grids by grasping only their edges for vitrification. This design also allowed the users to pick up the grids with consistent depth and angle, resulting in very consistent vitrification (**Fig. 1 g**). To finish the design, we also added a few more elements, such as a set of feet and a set of holes (**Fig. 1 b** and **c**). The users can then pick up the whole kit using a pair of tweezers easily without the kit being sucked on the bottom. We also noticed that when doing so, a layer of medium will be preserved between the cap and lid. Therefore, once cells are attached to the grids, the entire kit can be taken out and moved to another dish with new medium, for a fast, convenient and safe medium exchange (**Fig1. h**). This practice can be particularly useful when the users plate cells using a different condition than culturing cells.

**Fig 1.**
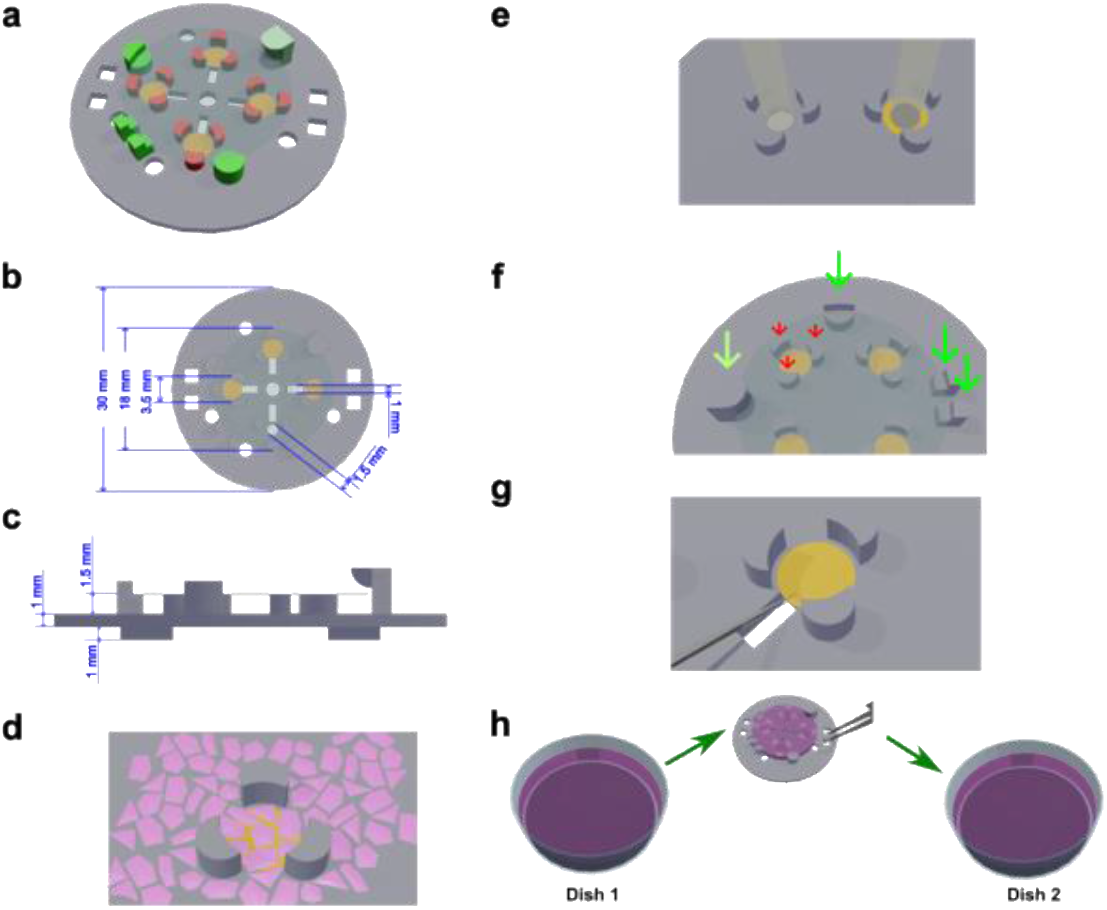
General design and usage of SLICK. **a**, Fully assembled SLICK loaded with 4 EM grids. The pillars that make up the grids slots are illustrated in red and the lock system for the lid (a coverslip) is illustrated in green. **b** and **c**, Dimensions of SLICK. **d**, The grid slot is composed of three separate pillars surrounding the grids, providing sufficient stability while still allowing access to the medium. **e**, Observation holes underneath the center of the grids enable real-time light microscopy monitoring. **f**, During experiment, grids are stabilized both horizontally by three pillars (red arrows) and vertically by a semi-locked lid (green arrows). **g**, A thin trench extends right underneath the edge of the EM grid, so the users can pick up the grid conveniently, safely and consistently. **h**, The entire kit can be picked up from the dish and moved to another dish for rapid medium change. In this process, small amount of medium will be preserved between the base and the lid, preventing the cells from drying.

We tested this kit by culturing rat hippocampus primary neurons for cryo-ET imaging. Neurons are highly differentiated cells and cryo-ET has been long considered a suitable tool to understand their ultrastructure^8^. However, preparing neurons for cryo-ET studies are typically difficult and sometimes yield suboptimal results^9^. Our test here is inspired by a well-established protocol where neurons are cultured on coverslips that are placed in dishes containing glial feeder cells^10^. With the feeder cells, neurons can be cultured at low density which is suitable for cryo-ET studies. For cryo-ET studies, we replaced glass coverslips in this protocol with SLICKs loaded with EM grids. After culturing 3-4 weeks, the SLICKs will be opened, and grids will be picked up for vitrification (**Extended Fig. 3**). Subsequently collected cryo-ET data showed well developed synapses, suggesting SLICK can be useful for preparing some of the most dedicated cells and compatible with very complicated culturing procedures (**Fig. 2**). Obviously, more robust cancer cells such as HeLa cells can also be easily cultured by SLICK with high throughput (**Extended Fig. 4**).

**Fig 2.**
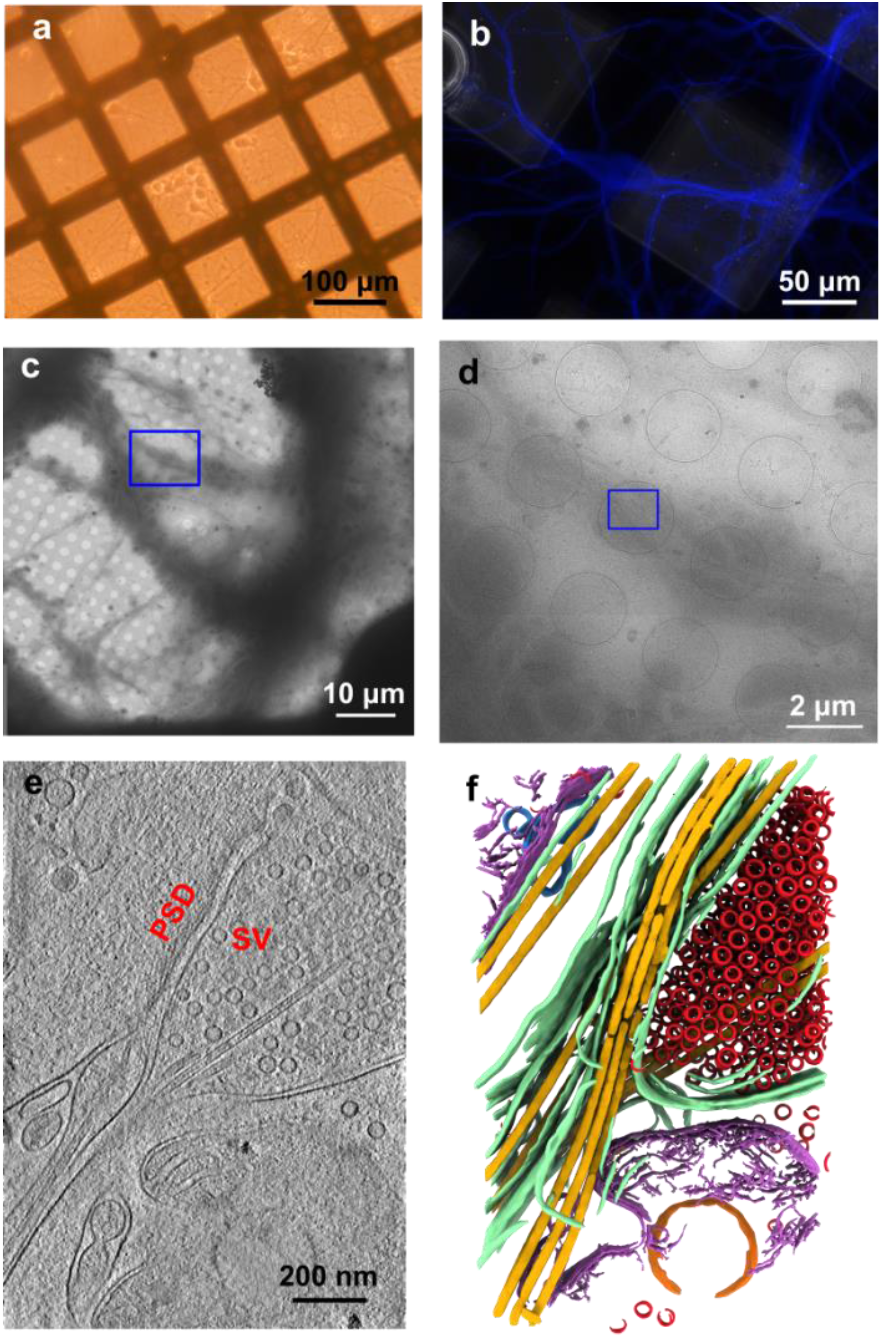
Testing SLICK by culturing low density of primary hippocampus neurons. **a**, Loaded SLICKs can be placed in dishes with feeder cells, therefore enabling low density neuron culture on EM grids. **b**, After three weeks of culturing, primary neurons have shown substantially developed dendrites as visualized by immunolabelling MAPT2. **c** and **d**, Neurites and synapses can be found under electron microscopy for further high-resolution imaging. **e**, One slice from a representative reconstructed tomogram showing synaptic vesicles (SV) and post-synaptic density (PSD). **f**, The representative tomogram in **e** was annotated to show different cellular component including: plasma membrane (green), microtubules (yellow), mitochondria (purple), synaptic vesicles (red), vesicles (orange), and endoplasmic reticulum (blue).

The above described and tested SLICK contains all the essential elements for high-throughput sample preparation. While previous attempts have utilized 3D-printed grid holders to facilitate cryo-ET sample preparation, these holders feature three walls that enclose the EM grids, restricting their accessibility to the surrounding microenvironment^11^. Additionally, the presence of these walls may create uneven horizontal material exchange, potentially altering physiological conditions for cells on the EM grid. In contrast, the SLICK system presented here demonstrates a more sophisticated design that improves grid stability while minimizing interference with the culturing environment. The three-pillar architecture allows cells on the EM grid to communicate with neighboring cells within the same microenvironment, preserving their physiological activity. SLICK is highly versatile and can be further customized for specific applications (**Extended Fig. 5**). For example, we frequently coat the SLICK base with approximately 10 µm of parylene to enhance biocompatibility, particularly for primary neuron cultures^12^. The size and grid capacity of SLICK can also be adjusted to meet user requirements. With SLICK, it is now feasible to culture cells on 12 EM grids using a single 35 mm dish. These grids can be directly loaded into modern FIB-SEM and TEM systems using a standard 12-slot EM cassette^13^, eliminating excessive steps of loading and unloading, thereby streamlining the workflow and significantly increasing the throughput of cryo-ET data collection. For applications requiring even higher throughput, SLICK can accommodate over 150 EM grids in a single 100 mm dish. This scalability is particularly advantageous for studying rare biological events or low-abundance macromolecules, which are challenging to capture with current throughput limitations.

Despite these advancements, there remain opportunities for further improvement. For instance, the current lid design is loosely placed on the base, requiring extra attention from the experimenter. Addressing such limitations will likely require more sophisticated manufacturing techniques and complex designs, which may be challenging to implement in a typical cell biology laboratory setting.

We anticipate that this kit will significantly improve the throughput and consistency for current *in situ* cryo-EM practitioners and potentially opens the windows for time resolved experiments. More importantly, we believe this kit will lower the technical barrier to *in situ* cryo-EM, enabling broader access to the state-of-the-art method (1379 words).

## Methods

### Design and Manufacturing of SLICK

3D models of SLICK base were all designed in Blender 4.3.1 (The Blender Foundation, Amsterdam, Netherlands). The 3D printing orientation was adjusted in PreForm 3.40.1 (formlabs, Massachusetts, US) to ensure that the supports were not placed on the top surface of the SLICK base. All SLICK bases were printed on a Form 2 printer using clear V4 resin (formlabs, Massachusetts, US). After removing the supports, SLICK bases were rinsed briefly in isopropanol and dried in air. SLICK bases were coated with 10-50 μm of parylene (coated by VSI Parylene, Colorado, US, or using Labcoater®-PDS 2010 equipment (Specialty Coating Systems, Indiana, USA) at Microfabrication Laboratory at Case Western Reserve University).

New coverslips were soaked in 1M HCl overnight and rinsed extensively with ddH2O, then sonicated in ddH2O and washed by ddH2O again for over 6 times. The coverslips were preserved in 70% ethanol until usage^14^.

### Assembly of SLICK

A SLICK base was placed within a 35-mm Petri dish (e.g., cat# 353001, Corning Inc. NY, USA), and gold EM grids (e.g., 200 mesh QUANTIFOIL R 2/1 carbon grids or 200 mesh QUANTIFOIL R1.2/20 SiO2 grids) were loaded into the grid slots. The entire assembly was glow discharged at 25 mA with negative polarity for 30 seconds in a PELCO easiGlow glow discharge unit (TedPella, California, USA). After glow discharge, the SLICK base was capped with a preserved #1.5 coverslip (18 mm round coverslip with 0.16∼0.19 mm in thickness). This fully assembled kit should be used immediately.

### Preparing SLICK for Plating

Four milliliters of 70% isopropanol (Fisher Scientific, USA) were added to the 35-mm dish containing fully assembled SLICK, and the dish was closed and transferred to a tissue culture hood. After 15 minutes, the SLICK was rinsed three times with sterile water. The kit was then incubated in 1 mg/ml poly-L-Lysine (ThermoFisher Scientific, MA, USA) in room temperature overnight for primary neurons, or incubated with 0.05 mg/mL Type I collagen (Gibco, A10644-01) in 37°C for 1 hour for HeLa cells. After incubation, the kit was rinsed with sterile water for 3 times and incubated with medium for further use.

When adding liquid, it is recommended to use a long pipette (> 5 ml) and to firmly press down the coverslip with the pipette tip for optimal stability. Additionally, sufficient liquid should always be added at once to fully immerse the coverslip to prevent it from floating.

### Plating and Culturing of Primary Neurons

The hippocampal neuron culturing procedure was adapted from a previously established method where coverslips were replaced by assembled SLICKS, with the following modifications. 1) glial feeder layers were prepared in 35-mm dishes; 2) SLICKs were cleaned and coated as mentioned above before equilibrated in Neuronal Plating Medium in a CO2 incubator at 37°C overnight; 3), approximately 500,000 dissociated hippocampal neurons were added to a 35-mm dish containing the SLICK and mixed gently; 4) after cell attachment was confirmed by light microscopy, the SLICKs were directly transferred to dishes with the glial feeder layer. Cells were maintained as previously described, and neuronal growth was monitored by light microscopy.

Several additional practices may improve neuronal growth, though they have not been extensively tested. For instance, rinsing the SLICK with Neuronal Maintenance Medium before transferring it to dishes with the glial feeder layer may help remove residual Neuronal Plating Medium. Additionally, the SLICK base and lid can be replaced with new ones during the transfer, or the EM grids can be flipped so that the neurons directly face the feeder layer.

### Immunostaining of primary neurons

After 3 weeks of culturing, primary neurons on EM grids were fixed in pre-warmed 4% Paraformaldehyde (PFA) containing 4% sucrose in PBS for 12 minutes. Neurons were washed three times with PBS, including one time with 0.2% TritonX-100 in between for 5 minutes, and incubated in a blocking solution (5% normal goat serum, 3% BSA in PBS) for 30 minutes at 37 °C. Then neurons were labelled by anti-MAP2 (chicken, SYSY, cat#188006, 1:6,000) at 4 °C overnight, followed by AMCA-goat-anti-Chicken IgY antibodies (Jackson ImmunoResearch, Cat# 103155155) in the blocking solution (37 °C 30 minutes). After three washes in PBS, grids were incubated in Elvanol medium. The fixing, washing, and blocking was done in a SLICK dish, while antibody labeling was performed by placing the grid on a drop of diluted antibody, with the cell side facing the droplet.

### Plating and Culturing of HeLa cells

HeLa cells, previously maintained at ∼40%-80% confluency, were trypsinized, centrifuged at 300 × g for 3 minutes, and resuspended in fresh maintenance medium (DMEM with 10% FBS and 1% penicillin/streptomycin) to a final density of 100,000 cells/ml. A 4-ml aliquot of the well-mixed cell suspension was added directly to a 35-mm dish containing SLICK. After overnight culturing, the EM grids were ready for vitrification.

### Confocal Imaging of HeLa Cells on EM Grids

HeLa cells were cultured overnight and stained with CellMask Plasma Membrane Staining Dye (Invitrogen C10046) in PBS for 5 minutes. After staining, the SLICK-containing EM grids with attached cells were washed with PBS and fixed with 4% paraformaldehyde for 20 minutes. Following fixation, the grids were thoroughly washed and immersed in PBS buffer. The EM grids were then transferred to a new 35 mm petri dish containing PBS buffer for confocal visualization using an Olympus FV 3000 confocal laser scanning microscope.

Montaged images were acquired using an auto-focus scheme, capturing 16 individual images to cover the entire grid area. The images were subsequently stitched together to generate a complete view of the grid.

### Vitrification and Lamella Preparation

Once cells reached the desired stage (3 weeks for primary neurons and overnight for HeLa cells in this study), the coverslip lids were removed, and the grids were manually blotted and plunge-frozen in liquid ethane and transferred into liquid nitrogen. HeLa cells on the clipped EM grids were milled by Aquilos 2 Cryo-FIB/SEM for obtaining a thin lamella with a thickness of 200 nm, as previously described^15^.

### Cryo-ET Data Collection

All tilt series were collected using SerialEM on a Titan Krios G3i electron microscopy equipped with a BioQuantum K3 direct electron detector operated at 300 KeV in zero-loss mode (slit width 20 eV). For HeLa cells lamella sample, the stage was pre-tilted by 9º, and the remaining of data collection were performed in the same manner for both cell types as described below. Tilt series were collected in dose symmetric scheme with a tilt range from -51º to 51º and an increment of 3º. 8 frames per tilt were collected with a physical pixel size of 0.25 nm. Defocus was set between 6-8 μm and the total electron dose was set to ∼ 100 e^-^/Å^2^.

### Cryo-ET Data Processing

Frames were aligned and summed in Warp^16^ 1.0.9, then exported as unaligned tilt series. Tilt series alignment and tomogram reconstruction were carried out in AreTomo^17^ 1.3.4 using weighted back projection (WBP) with a binning factor of 4. Reconstructed tomograms were denoised with TomoAND^18^ and annotated in EMAN2 (version 2.99.47) using the convolutional neural network method^19^.

## Supporting information

Stability test for SLICK

## Acknowledgements

We thank K. Li and K. Whiddon in CCMSB cryo-EM core for the assistance with cryo-ET data collection

## Funding

We acknowledge grants support from the NIH National Institute of Mental Health (1R01MH130476) and Simons Foundation Autism Research Initiative Bridge to Independence Award to P.Z; Case Western Reserve University start-up funds to S.Z.

## Author contributions

Q.L. performed the experiments and wrote the paper. L.Z. performed cryo-FIB/SEM experiments and confocal microscopy observation. Q.X. and P.Z. performed primary neuron culture on SLICK alongside Q.L. S.Z. conceptualized the project and wrote the paper.

**Extended Fig 1.**
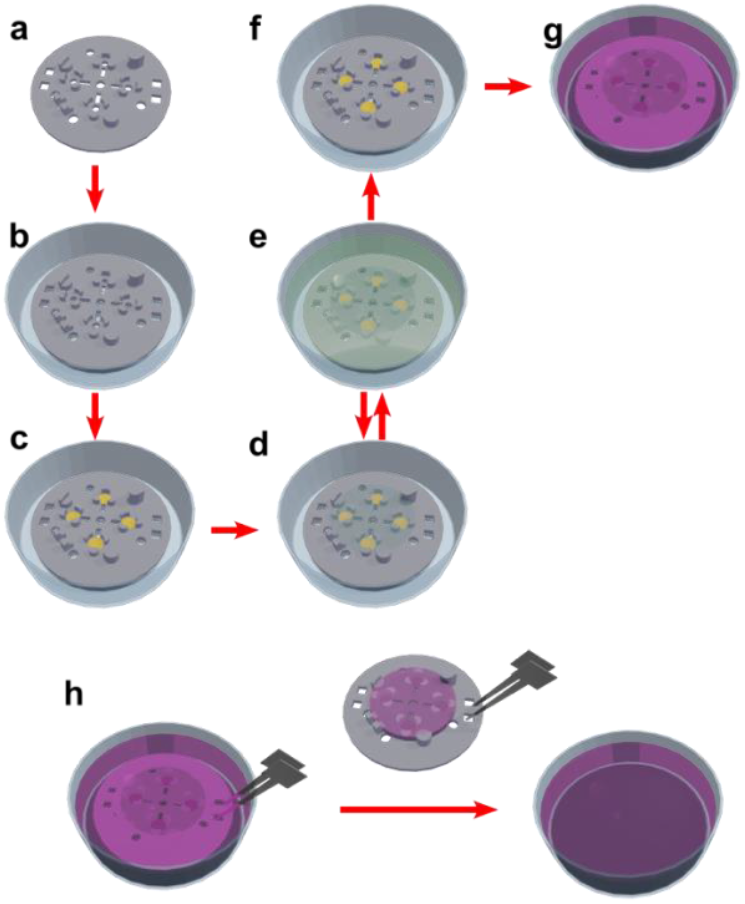
Typical experimental procedure for preparing adhesive cells using SLICK. Properly cleaned SLICK base (**a**) will be first placed in a Petri dish (**b**). The EM grids will be loaded into the SLICK base for glow discharge or other surface treatment (**c**). After this step, the lid (a coverslip) will be closed till vitrification. Such an assembly can stably go through repeated sterilization, coating, and washing (**d** and **e**) with minimal chance of damage. When the system is ready for plating (**f**), cell suspension will be added to the Petri dish (**g**). If medium change is needed, the entire SLICK can be conveniently moved to a new dish containing the new medium (**h**).

**Extended Fig 2.**
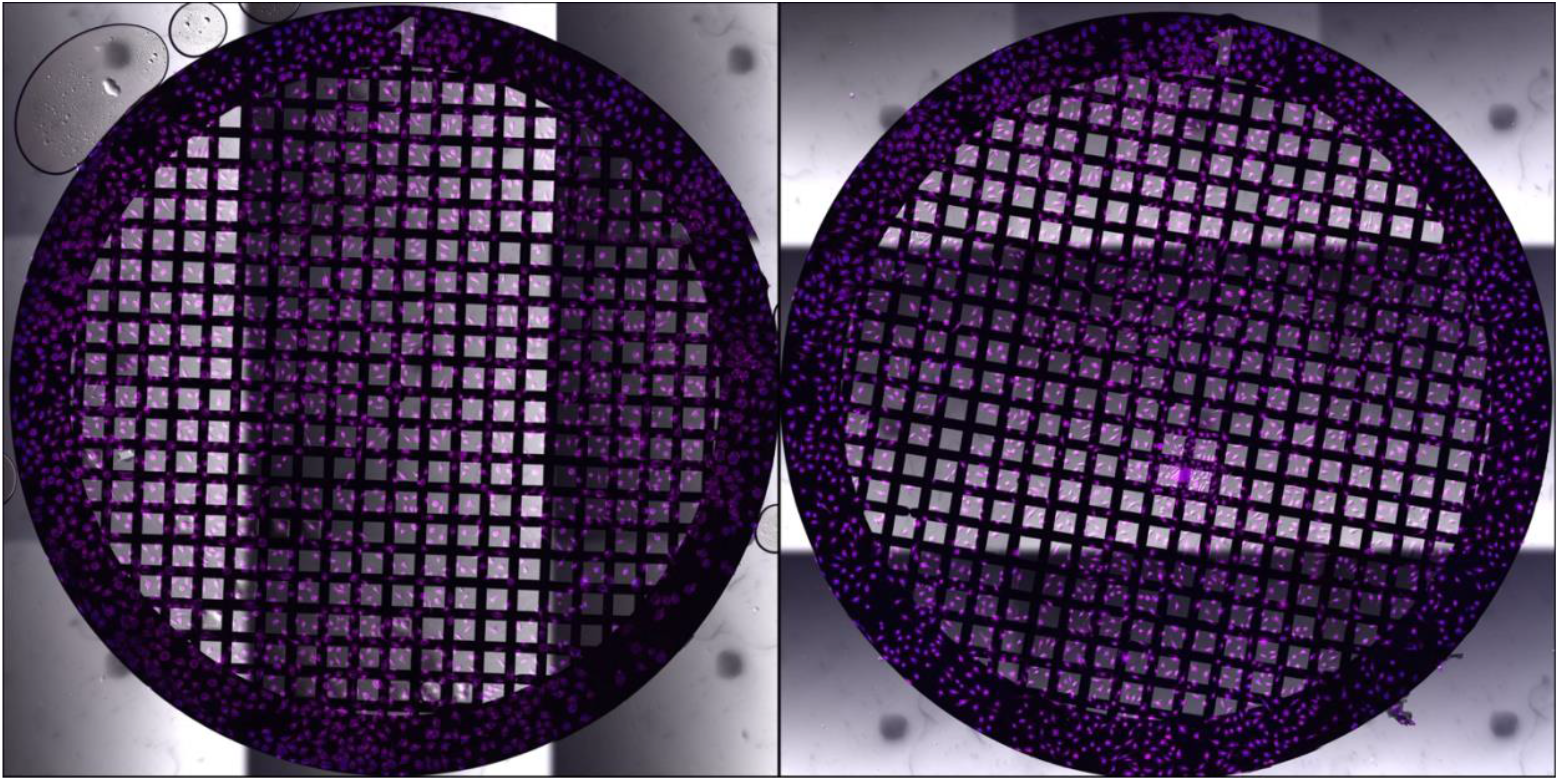
Representative light microscopy images of EM grids after overnight culturing. Each grid was only handled by tweezers twice: before glow discharge and during plunge freezing. Therefore, the chance of grid damage is minimized.

**Extended Fig 3.**
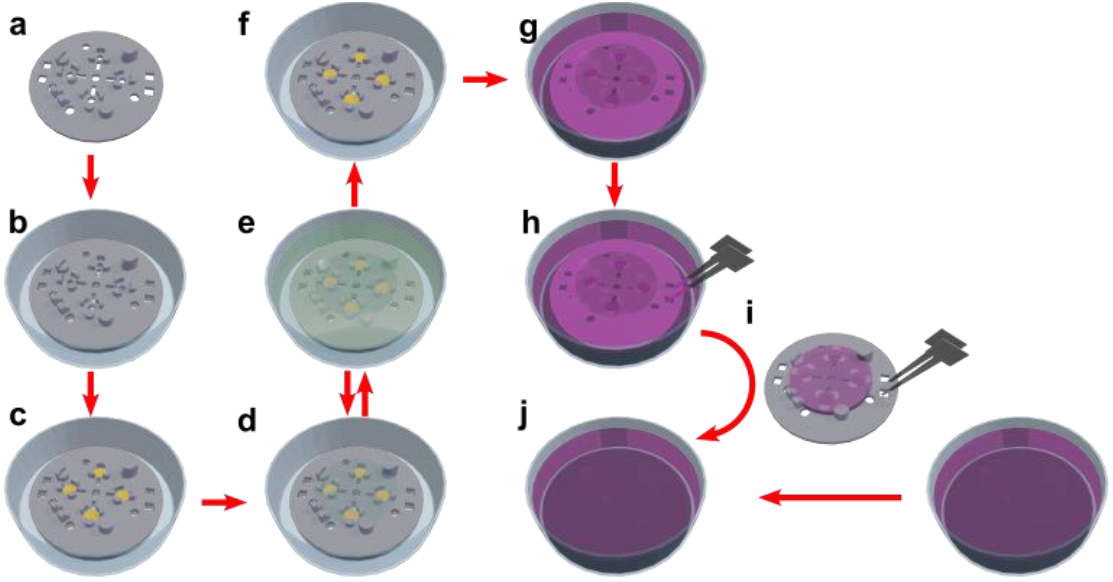
Experimental procedure for preparing low density primary neurons with feeder layers using SLICK. Properly cleaned SLICK base (**a**) will be first placed in a Petri dish (**b**). The EM grids will be loaded into the SLICK base for glow discharge or other surface treatment (**c**). After this step, the lid (a coverslip) will be closed till vitrification. Such an assembly can stably go through repeated sterilization, coating, washing, and equilibrium (**d** and **e**) with minimal chance of damage. When the system is ready for plating (**f**), primary neuron suspension will be added to the Petri dish (**g**). After neurons are attached to the EM grids, SLICK will be moved to a dish with feeder cells for culturing.

**Extended Fig 4.**
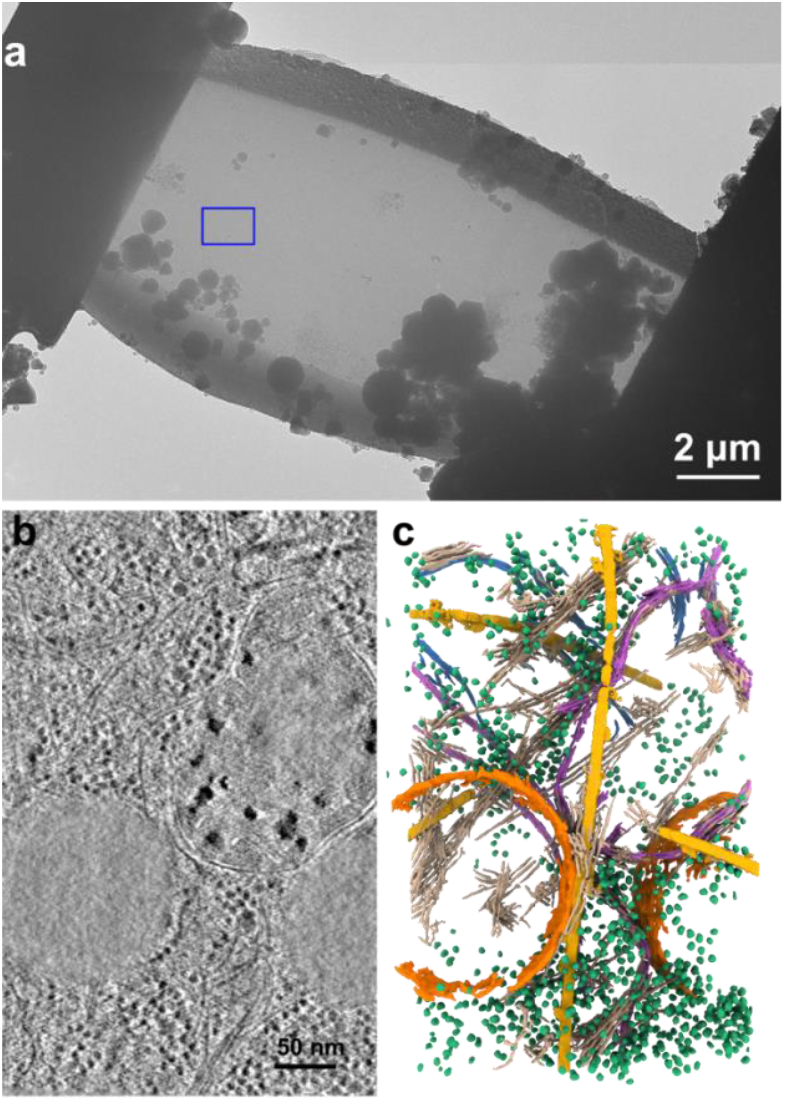
SLICK can also be used to prepare thicker cells for focused ion beam (FIB) milling and subsequent cryo-ET studies. **a**, A representative lamella prepared from HeLa cells prepared using SLICK. **b**, One slice from a representative tomogram collected from the lamella in **a. c**, Annotated tomogram in **b**, with ribosomes in green, vesicles in orange, microtubules in yellow, septins in brown, mitochondrion in purple, and endoplasmic reticulum in blue.

**Extended Fig 5.**
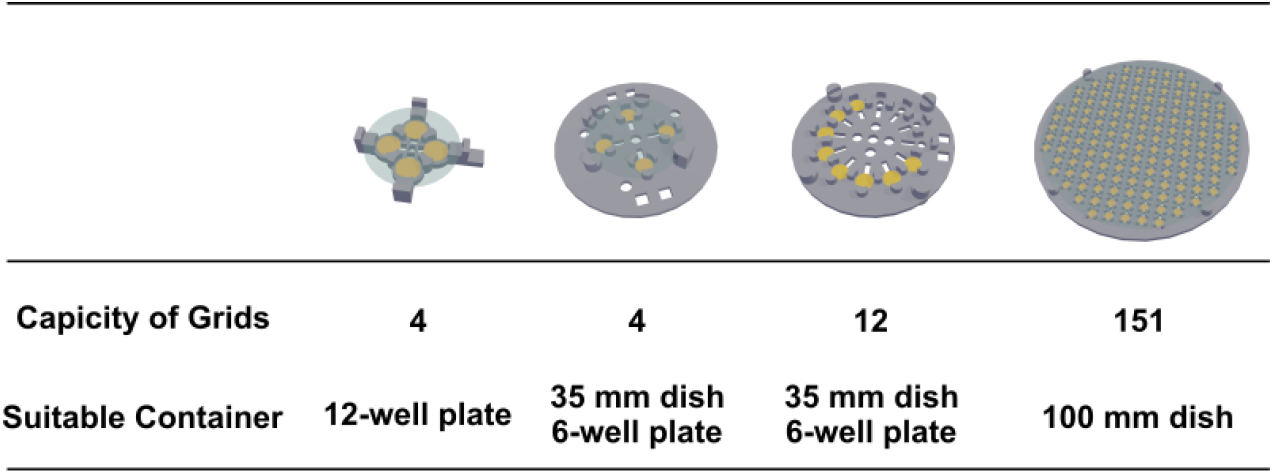
Alternative designs of SLICK for different culturing containers and different throughput.

**Supplementary Video 1. Stability test for SLICK**. A fully assembled SLICK is placed in a 35 mm dish filled with water, and the dish is placed in a shaker at 150 rpm. To facilitate visualization, the edge of the coverslip is colored in blue. Both grids and coverslip lid are secured in place during shaking.

## Notes

### Competing Interest Statement

The authors have declared no competing interest.

